# Biom3d, a modular framework to host and develop 3D segmentation methods

**DOI:** 10.1101/2024.07.25.604800

**Authors:** Guillaume Mougeot, Sami Safarbati, Hervé Alégot, Pierre Pouchin, Nadine Field, Sébastien Almagro, Émilie Pery, Aline Probst, Christophe Tatout, David E. Evans, Katja Graumann, Frédéric Chausse, Sophie Desset

## Abstract

U-Net is a convolutional neural network model developed in 2015 and has proven to be one of the most inspiring deep-learning models for image segmentation. Numerous U-Net-based applications have since emerged, constituting a heterogeneous set of tools that illustrate the current reproducibility crisis in the deep-learning field and that remain slow to spread in application fields such as 3D bioimaging. Here we propose a solution named Biom3d, a modular deep learning framework facilitating the integration and development of novel models, metrics, or training schemes for 3D image segmentation. The development philosophy of Biom3d provides an improved code sustainability and reproducibility in line with the FAIR principles and is available as a graphical user interface and an open-source deep-learning framework to target a large community of users, from end users to deep learning developers.

## Introduction

The biomedical field produces many three-dimensional images (3D), whether to follow the fate of a tumor in an organ or to measure shape properties of cells, nuclei or other organelles. Such properties can be extracted using a method in image analysis called semantic segmentation. Semantic segmentation classifies each pixel from an image as belonging to the object of interest or to the background and thus produces a binary mask of the image. The increased massification of data has led to the development of innovative solutions to automate segmentation. While these solutions present some challenges, especially when it comes to 3D biomedical images, they offer great potential. The variability in imaging modalities, noise-to-signal ratio, size, voxel spacing or volume distribution of the objects to segment presents exciting opportunities for further development. This is where deep learning methods come into play, being both performant and flexible on a broad range of applications, from denoising^1^ to image segmentation^2^, and of different modalities and scales^3^, from microscopy to medical imaging, both for 2D and 3D images.

U-Net, a convolutional neural network model developed in 2015^4^, has been demonstrated to be one of the most effective deep-learning models for 3D semantic segmentation. Numerous U-Net-based applications^2^ have since emerged but are still not routinely used for bioimage analysis. This can be attributed, in part, to the difficulty encountered by non-programmers in mastering such tools. Furthermore, there is a lack of evidence to demonstrate the superiority of deep learning methods in comparison with conventional ones. Some methods, such as CellPose^5,6^, StarDist^7^ and DeepImageJ^8^, are ready to use, with the necessary code, documentation, and information about the computing environment^2^. Unfortunately, as they have become more specialized, they have lost a certain flexibility that limits their application to a wide range of images. Their possibility of new training, especially for 3D images, is not always straightforward or even possible. On one hand, nnU-Net^9^ exemplifies this flexibility by having an excellent ability to self-configure from training sets that can be quite different, adapting to many image acquisition modalities with good results, but remains difficult to use for users without programming experience. At the other end of the skill spectrum, deep learning developers have at their disposal a bank of functions perfectly documented and updated thanks to MONAI^10^, but they do not have an assembly plan allowing them to build a complete workflow such as proposed for 2D images by OpenMMLab^11^. The current situation is that the development of innovations in image analysis using deep learning remains a time-consuming process, often with relatively inflexible applications, and there is no suitable methodology for rapidly evaluating them. To fill these gaps, we have developed a deep learning segmentation framework that combines the ease of use of DeepImageJ^8^ with the flexibility of nnU-Net^9^, while also serving as a development base for improving methods for developers in the style of OpenMMLab^11^.

In this work, in addition to performance, flexibility and reusability, we propose a new prism through which to evaluate a deep learning method: **sustainability**. Our definition of method sustainability encapsulates two main objectives: method accessibility and adaptability. Method accessibility relates to the range of users who will utilize the tool. Increasing this range would not only result in an enlarged user community, but would also facilitate the emergence of multidisciplinary interactions, a goal that is supported by a significant number of researchers^12^. This is particularly true for software inherited from fundamental research, which attempts to reach applied research. Involving end-users will ensure that the software meets the needs of an external community and has concrete applications, giving it both meaning and feedback for improvements. Involving external developers will attract novel ideas while strengthening a community of maintainers. Method adaptability relates to this last community of developers and is the ability of a tool to quickly evolve, especially in the fast-moving world of deep learning. It can be reasonably assumed that the more adaptable a tool is, and the broader its user community, the longer it should last.

Following this, we defined a new tool development philosophy, emphasizing audience broadening, code clarity and code modularity for deep learning applications. To reach a wider audience, our aim is to create a series of tools that bridge the gap between end-users and developers in a gentle way, meaning that end-users wishing to deepen their understanding of the method right down to the lines of code need to be able to do this in small steps only. For each user profile, the objective is thus to substantially reduce the time required to invest in using a deep learning method, while simultaneously expanding the scope of accessible functions (Figure 1). The intention is to prioritize usability to the same extent as functionality. Current state-of-the-art methods usually target only one of these user profiles: nnU-Net^9^, Cellpose^6^ or StarDist^7^ target Python Programmers, while ZeroCostDL4Mic^13^ or DeepImageJ^8^ are mainly addressed to Non-Programmers (Figure 1). Moreover, during the code implementation, the emphasis should be made on code clarity, which implies that the code should remain comprehensible from its overall architecture to its individual lines by, for example, being presented in a hierarchical structure. This approach starts with high-level, general functions and progressively delves into more granular details and functionalities, facilitating a gradual and systematic comprehension of the code.

**Figure 1.**
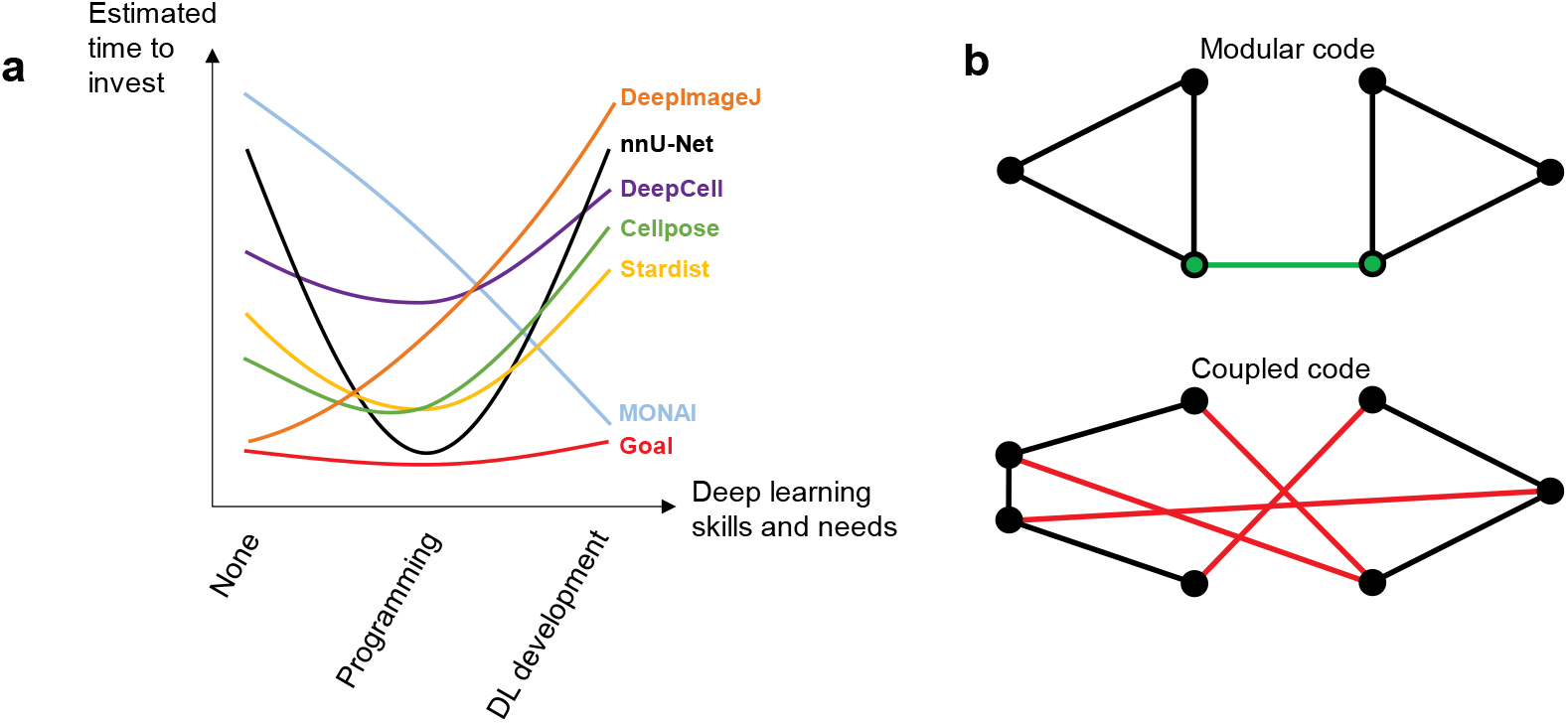
Method accessibility for different user profiles. **a**, Fulfilling the needs of a continuum of user profiles. In the abscissa are conjointly represented the skills and needs of deep learning users while in the ordinate is represented the estimated time required for them to use the method and fulfil their needs. On the left, non-programmers may expect easy-to-use software with graphical user interfaces and better segmentation results than with non-deep learning methods, without the need for delving into deep learning theory. In the middle, Python programmers may look for a command line interface or a Python package as well as a performant, flexible and easily reconfigurable deep learning tool. On the right, deep learning developers may be interested in a well-documented and modular deep learning framework, where pieces of code can easily be changed, removed, or added. Arbitrary lines represent our experiences to install, use, modify these methods in our laboratories. Most of the state-of-the-art methods are only targeting one user profile, noticeably omitting deep learning developers. The Goal line can be defined as the ideal method targeting all profiles. **b**, Difference between a modular code and a coupled code. Each piece of code represents a node in these graphs. Each edge is an interaction between two distinct pieces of code. A modular code (*top*) means that the few code components are clearly isolated and have few clearly defined links. A coupled code (*bottom*) has code components with a low internal cohesion and are strongly intertwined.

Such a method can also be adapted to novel innovations by incorporating an additional objective into the development process: code modularity. Code modularity means that components should be as self-contained and independent as possible (Figure 1). In computer science, this concept is also known as code cohesion and is opposed to code coupling^14^. This approach allows programmers to seamlessly incorporate or remove specific code components without causing undue disruptions to the overall system.

Following all these sustainability constraints, we developed a series of several open-access tools called Biom3d. By default, it is optimized to segment 3D images potentially having multiple channels or simultaneously presenting objects from different classes. These volumetric images can originate from CT-scan, MRI, confocal, X-ray or electron microscope. Biom3d integrates offline and online interfaces (GUI, CLI, and API), automated configuration processes, editable configuration files and a modular deep learning framework facilitating the choice and implementation of new models, metrics, data loaders and training routines.

## Results

### Biom3d flexibility and performance

Biom3d was first benchmarked with images from different modalities (CT-scan and MRI) including several classes of segmented objects (1 to 13). We chose well-known medical datasets from the Medical Segmentation Decathlon (MSD)^15^ and the Multi-organ Abdominal challenge (often called BTCV)^16^ that were used to develop the flexibility of nnU-Net (Figure 2a-d). To further assess its capabilities, Biom3d was also evaluated with several microscopy modalities on biological samples: X-ray (Figure 2e), electron microscopy (Figure 2f-g), and fluorescence microscopy (Figure 2h-j). Biom3d automatically adjusts its training configuration, resulting in Dice scores on the test sets reaching or exceeding those of nnU-Net (Figure 3a). For electron microscopy (EM) images, Biom3d achieved a Dice score of 92% when trained and evaluated with the two 3D images from the EPFL public database^17^ (Figure 3a). The Biom3d preprocessing workflow overcomes a limitation of nnU-Net by enabling the use of a single image for training, achieved by splitting the image into two (Supplementary Figure 5). Finally, Biom3d was applied to segment nonconvex objects such as X-ray images of aortic lamellae (Figure 2e) or as membranes of epithelial cells (Figure 2j). For all these cases, Biom3d successfully segmented the objects of interest (Figure 2).

**Figure 2.**
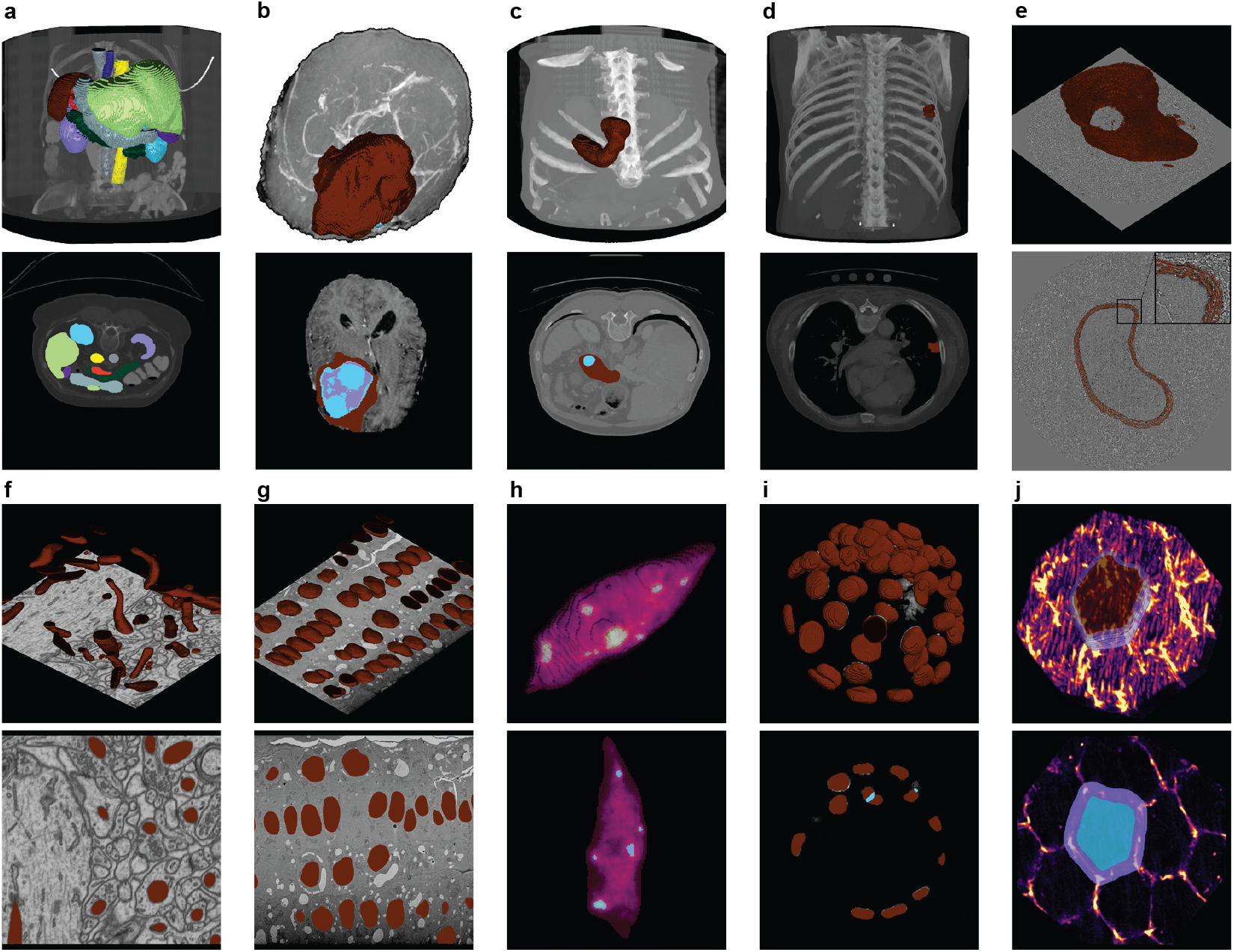
Samples of Biom3d predictions over several modalities from medical datasets (*top row*) and biological datasets (*bottom row*). All views originate from testing images, unseen during the training of Biom3d default model. **a-d** and **f** originate from public datasets while the rest are custom datasets. For each example, a 3D view is depicted on top of a 2D z-slice, both representing an overlay of the segmentation result and the raw image. The rendering was done by Napari (https://napari.org/). **a**, Multi-organ abdominal challenge (BTCV) representing thirteen segmented organs in CT-scan. **b**, Three brain tumor sub-regions in MRI. **c**, A pancreas (red) and a tumor (blue) in CT-scan. **d**, A lung tumor in CT-scan. **e**, An aortic lamella in a synchrotron microscope. **f**, EPFL dataset of mitochondria in an electron microscope. **g**, Plant cell nuclei in an electron microscope. **h**, A plant cell nucleus and its chromocenters in a structured illumination microscope. **i**, Nuclei of a mouse embryo in a confocal microscope. To help the model separate nuclei clusters, frontier regions between nuclei have been segmented (blue). **j**, An epithelial cell of a Drosophila embryo. Both the inner cell and cell membranes have been segmented. Complete datasets are available online (Supplementary Table 1).

**Figure 3.**
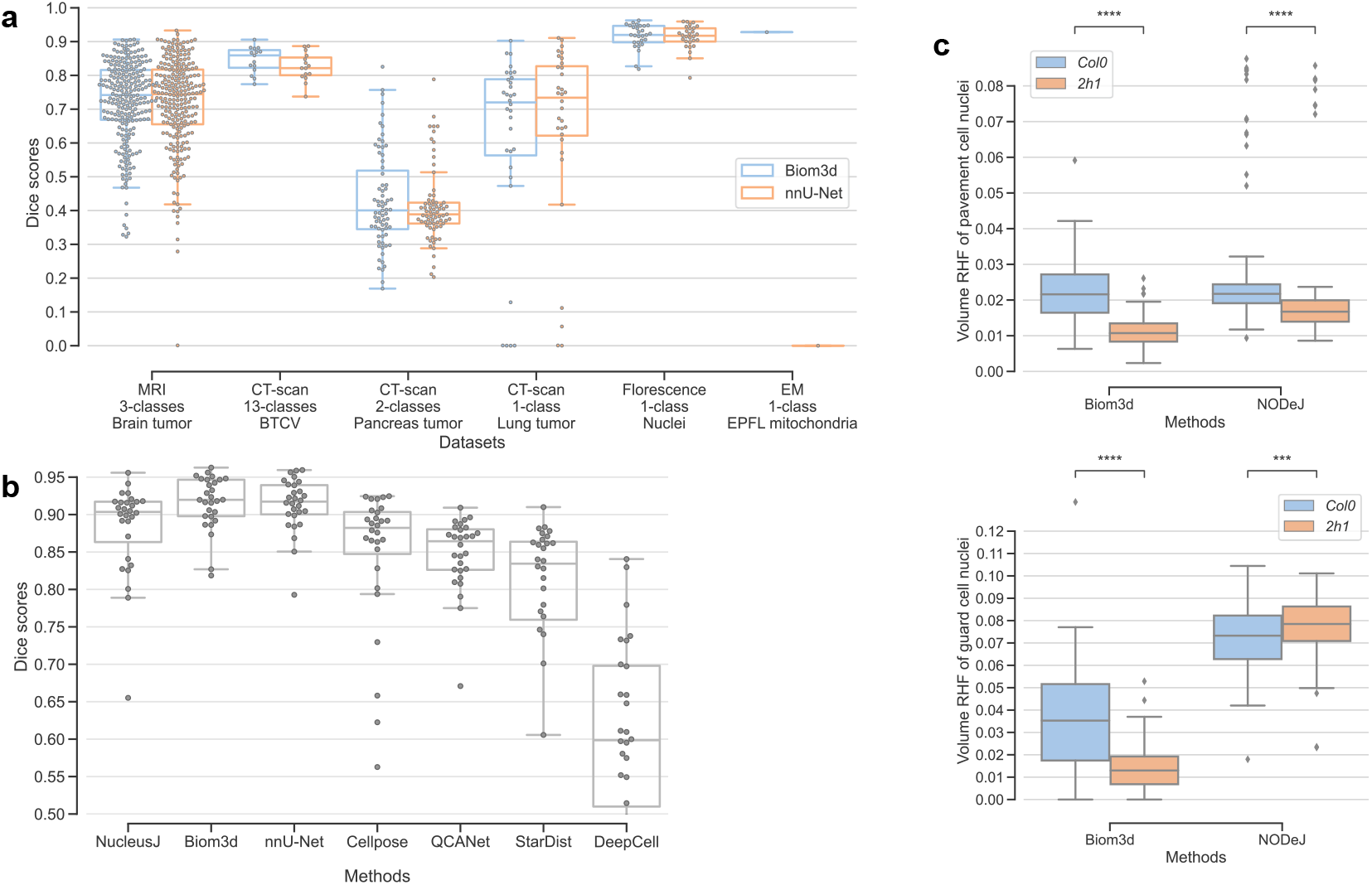
Biom3d performances. **a**, Biom3d achieves similar Dice scores to nnU-Net^9^ across various 3D segmentation datasets, which include different modalities and numbers of object classes. The Dice score, when applied to 3D segmentations, measures similarity by calculating twice the volume of intersection divided by the total volume of both the ground truth mask and the prediction. The higher the Dice score is, the better. The images in these datasets were captured with CT-scan (BTCV, Lung tumor, Pancreas tumor), MRI (Brain tumor), fluorescent microscope (Nuclei), and electron microscope (EPFL mitochondria). Public datasets from medical segmentation challenges (BTCV, Brain tumor, Lung tumor, and Pancreas tumor) were selected for their diversity in the number of object classes to segment. These datasets were split in two, with the first half of the images and masks used for training and the second half for testing. For the custom dataset Nuclei, 65 images and masks were used for training, while the remaining 28 were used for testing. The EPFL dataset has only one publicly available training image and one testing image. Training and testing images were identically chosen for Biom3d and nnU-Net. **b**, Benchmark of 7 segmentation methods on the Nuclei dataset, representing individual 3D plant nuclei. NucleusJ^18^ is a non-deep learning method specialized in 3D nucleus segmentation. Only Biom3d, nnU-Net and StarDist^7^ provided the requirement to train a new 3D segmentation model. Without a way to retrain them for 3D images, Cellpose^5^, QCANet^20^ and DeepCell^19^ were tested with their available pretrained models. Only Biom3d and nnU-Net exceeded the average Dice score obtained with NucleusJ. **c**, Example of Biom3d application to obtain biological insights. The volume of relative heterochromatin fraction (RHF) refers to the ratio between the volume of voxels located in chromocenter regions and of voxels located in the whole nucleus. The box plots represent samples of *A. thaliana* epithelial nuclei sorted between pavement cells (PC, *top*) and guard cells (GC, *bottom*). Biom3d is compared to NODeJ for segmenting chromocenters in nuclei from two plant types: *2h1* mutant^22^ and wild type. NODeJ, a non-deep-learning ImageJ plugin, was specifically developed to segment chromocenters in pavement cells (PC) using an intensity gradient method. While both methods produced expected results in PC nuclei, showing a strong decrease in relative heterochromatin fraction (RHF) in mutants, only Biom3d predicted a shift in RHF distribution in guard cell (GC) nuclei. NODeJ struggled to accurately segment regions of varying intensity in these smaller nuclei. The validity of Biom3d’s prediction was manually verified by experts.

To measure its performance against conventional methods, Biom3d was compared with NucleusJ^18^, specialized in 3D segmentation of individual plant nuclei. A custom dataset of 93 images of individual plant nuclei of various shapes and fluorescence intensity captured with a structured illumination microscope was created. In addition to nnU-Net and NucleusJ, Biom3d could also be compared to methods designed for 3D nucleus segmentation in fluorescent images^2^ like DeepCell^19^, QCANet^20^, Cellpose^6^ and StarDist^7^ (Figure 3b). While being slightly better than nnU-Net, Biom3d significantly outperforms the other methods, even the conventional method NucleusJ. Our hypothesis is that the pretraining of Cellpose, DeepCell and QCANet and the star-convex polyhedron constraint of StarDist introduce strong biases and prevent the model from appropriately fitting to the strong variation of intensity within the nuclei.

To investigate the limits of segmentation methods for answering a biological question, we finally compared Biom3d with NODeJ^21^, a conventional ImageJ plugin dedicated to segment high intensity spots into DAPI-stained nuclei named chromocenters. A custom dataset of 85 images of individual wild type plant nuclei captured with a structured illumination microscope was created to train Biom3d. The trained model was applied on a dataset of 698 images from wild type or 2h1 mutant plants. 2h1 is a mutant that has previously been reported to exhibit chromocenter loss^22^. While both methods correctly predicted chromocenter reduction in the nuclei of large pavement cells (PCs), only Biom3d was able to predict this expected result in small guard cells (GCs) (Figure 3c).

### A modular architecture

During Biom3d development, a significant amount of effort was dedicated to ensuring code modularity. It would be beneficial for programmers to have the opportunity to experiment with a broad degree of freedom in the deep learning hyper-parameters by altering the type of deep learning model, the learning rate, or the data loading process. These challenges are tackled with the modular structure of Biom3d, which first core components are Configuration Files. The Configuration Files of Biom3d depart from those of OpenMMLab being both simplified and completed. They include the definition of two types of hyper-parameters. First, stand-alone hyper-parameters can be integers, float numbers, or lists. They include parameters defined in the Graphical User Interface (patch size, number of epochs etc.) as well as many additional stand-alone hyper-parameters such as the initial learning rate or whether to use half-precision float-point format or not. Second, module hyper-parameters are key-value dictionaries. In this dictionary, the first key-value pair precises the name of the module being used, and the second key-value pair defines the parameters of this module. For example, “UNet3DVGGDeep” is the name of the 3D U-Net model in Biom3d, and its set of parameters is the number of pooling layers of the model and the number of classes of objects in the images (Figure 4a).

**Figure 4.**
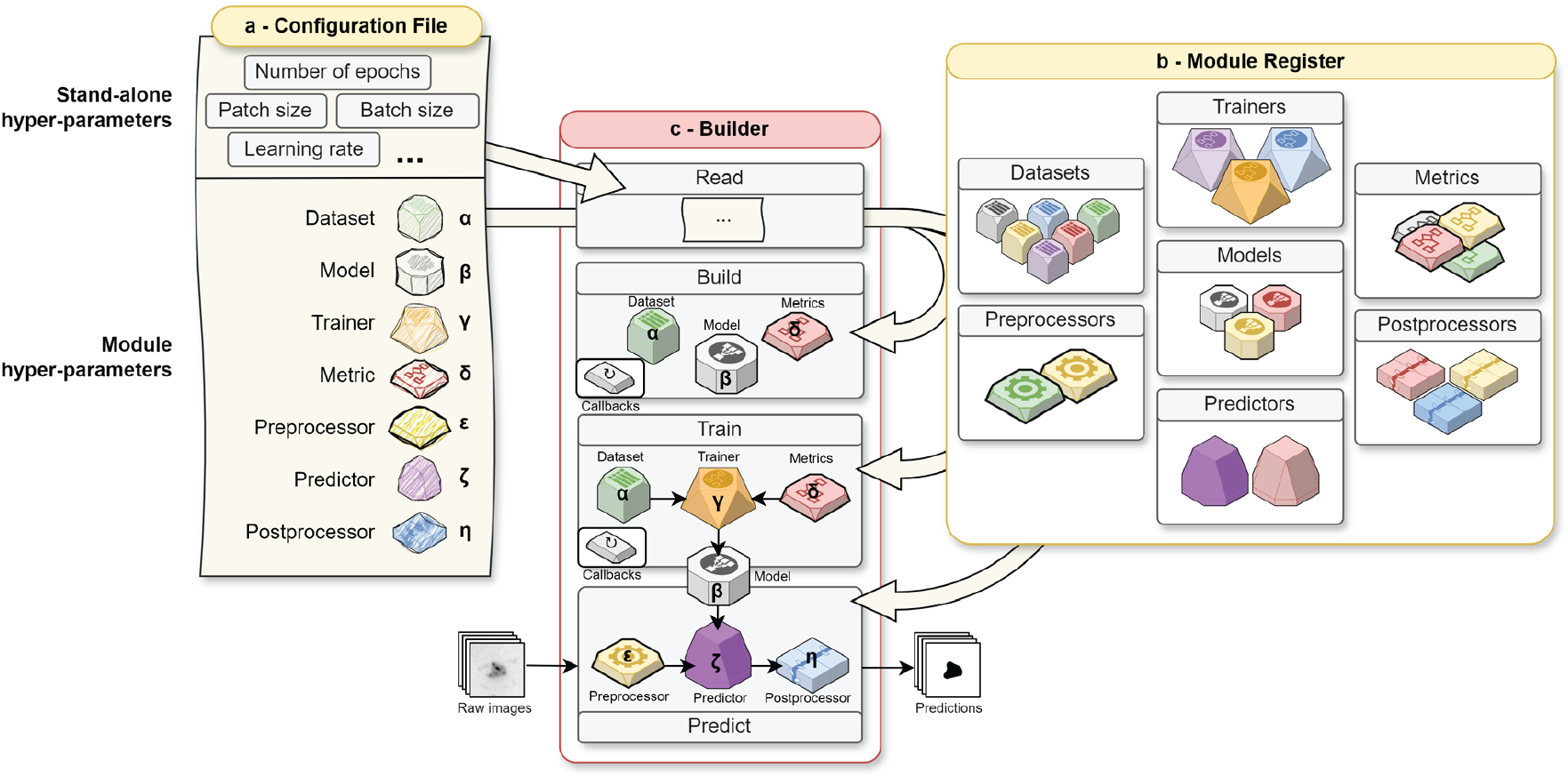
Biom3d workflow – Configuration File, Builder and Module Register. **a**, A Configuration File is an assembly plan defining Biom3d workflow for building, training and predicting with a deep learning model. It includes stand-alone hyper-parameters, such as the number of epoch or the learning rate, and module hyper-parameters, which represent the major building blocks of the workflow. A module hyper-parameter is defined with a name of an existing Module appearing in the Module Register and with its parameters (*Greek letters)*. **b**, The Module Register lists all existing modules within Biom3d, except for Callbacks which are defined in the Builder. Currently, there are seven different types of modules in the Module Register, each with multiple variants (*color shades*). **c**, The Builder reads all hyper-parameters from the Configuration File, retrieves the corresponding modules from the Module Register and builds them with their associated parameters. The Builder can then be used to train a Model with a Dataset, a Trainer, and a Metric. Once trained, the Model can be utilized by the Builder to predict segmentation masks from raw images by employing a Preprocessor, a Predictor, and a Postprocessor.

**Figure 5.**
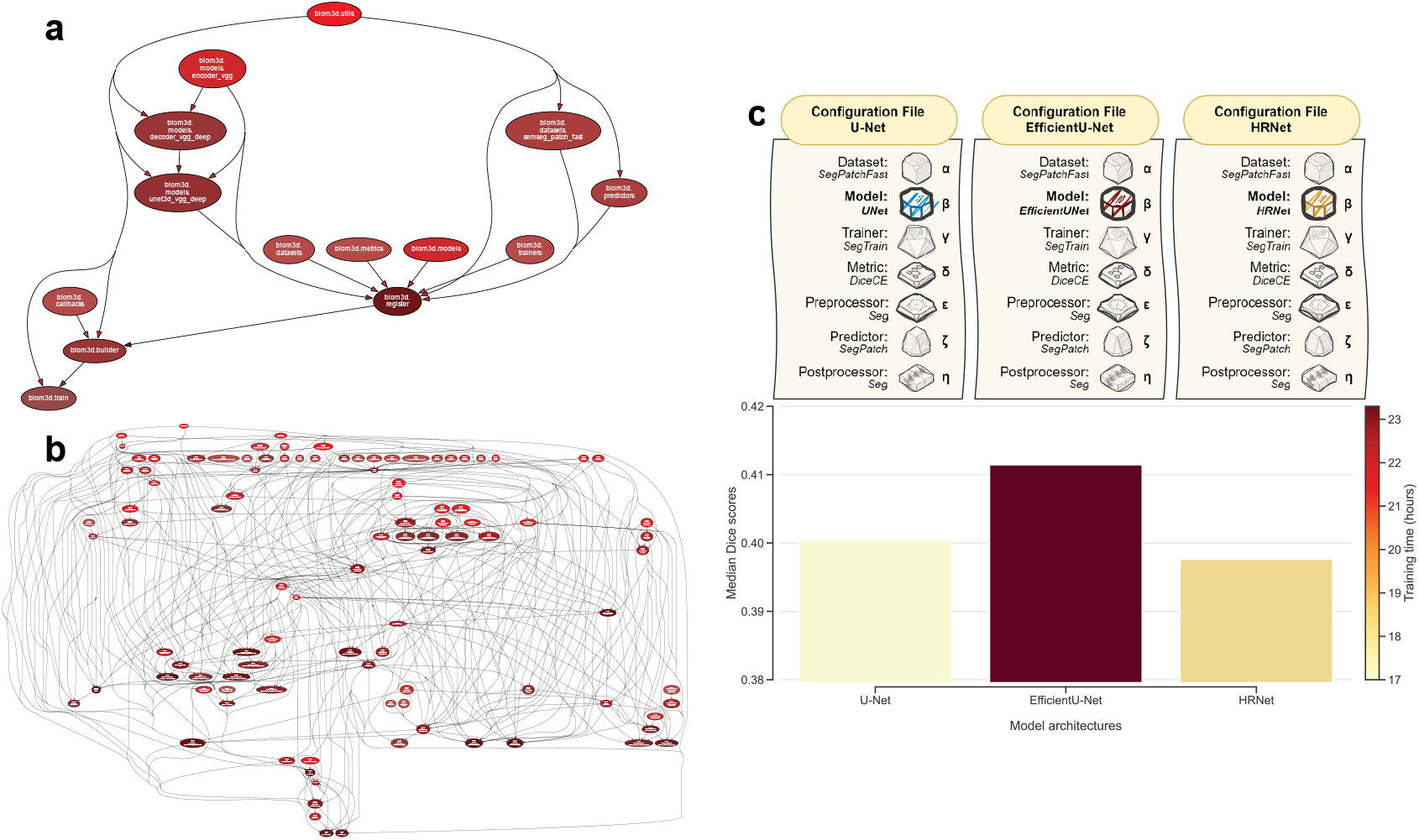
Code modularity of Biom3d. **a**, Representation of Biom3d code architecture. Each node in this directed acyclic graph is one Python script in Biom3d. Each directed edge is one call of one script by another. This graph was automatically generated by *pydeps*, a Python dependency visualizer, and by calling the default training workflow of Biom3d (*bottom left node*). The training process instantiates a Builder which then consults the Register to instantiate requested modules from the Configuration file. **b**, Representation of nnU-Net code architecture. For comparison with Biom3d, this graph was also generated by *pydeps* and depicts the nnU-Net training workflow. This graph indicates that nnU-Net code is more coupled than Biom3d code and that nnU-Net code will probably be harder to understand or modify. **c**, Exploiting modularity to benchmark model architectures. Once new model architectures have been added to Biom3d, a benchmark experiment can be conducted by only changing the model’s name in the configuration files, while leaving the rest of the training hyper-parameters untouched. The default U-Net model in Biom3d is here compared to an EfficientU-Net and a HRNet, both being integrated into Biom3d. These model architectures are trained and tested on the Pancreas tumor dataset. The results of this experiment indicate that, despite being slower, the EfficientU-Net model is more appropriate for this dataset than the others.

The second modularity component of Biom3d is the Module Register. Modules are code components classified in seven categories (Figure 4b): Preprocessors, preparing the datasets before training, Datasets, loading the datasets during training, Models, defining the deep learning models, Metrics, defining the loss functions for model performance assessment, Trainers, defining the training and validation loops, Predictors, defining the prediction loops, and Postprocessors, processing the model output before saving. All available Modules in Biom3d are named and listed in the Module Register. We expect that such a paradigm will simulate a significant number of deep learning methods, with a broader application spectrum than just image segmentation. While editing the Configuration File, a user can thus select from the Register the modules that fit the most to the expected task. Biom3d also includes an extra, special, eighth type of Module, Callbacks (Supplementary Figure 7), which decouple all recurrent events from the training loop, such as model saving, learning rate updates, logs printing and saving, or loss updates. Because the way Callbacks are depending on numerous hyper-parameters, their code definition is currently located directly in the Builder. As modules are Python classes or functions, if the user does not find the appropriate modules, new modules can be easily integrated into Biom3d by importing them and adding a new entry to the Register. It must be noticed that Biom3d accepts Pytorch modules from any origin if they respect the expected inputs and outputs of the surrounding modules. For example, if intending to add a new Metric to the default Biom3d configuration, the user must only make sure that it includes a few arguments like a name, a value and an average value. Once added to the Register, the new Module can directly be incorporated in Biom3d workflow by simply editing the Configuration File.

If Modules are the organs of Biom3d, the Builder is its skeleton, providing a general structure for the Module to properly work together (Figure 4c). The Builder reads the Configuration File, retrieves and instantiates the selected modules from the Register, and prepares them for training or prediction. Once a Builder is instantiated with a Configuration File, the user can then easily start the training or prediction process by calling a single function. The Builder stores its configuration and the training status (training curves, etc.) which can then be reused to carry on interrupted training or to fine-tune models with a new Configuration File.

### An easy-to-use tool

Biom3d integrates one local Graphical User Interface (GUI) and one online GUI in the form of a Google Colab notebook. An important innovation with Biom3d interfaces is the “auto-configuration” button. Once the dataset folders path has been entered and this button pressed, Biom3d proceeds to a cascade of hidden operations (Supplementary Figure 1). The dataset is first scanned to check its quality and to retrieve its characteristics, such as the median image size or the median sampling. These data characteristics are then used to normalize the image calibration and intensity and to automatically configure the training process of the deep learning model (Supplementary Figure 4). Biom3d is flexible on the input image format, as it works with most medical formats (Nifti, DICOM etc.) and TIFF format, largely used for microscopy data. Biom3d data scanning process is equipped with many safeguards to automatically correct most user annotator mistakes, which seems to be decisive in avoiding the “first-click abandonment” phenomenon.

The result of this automated training configuration, including parameters such as patch size and batch size, is stored in a YAML file. This file is partially displayed in the local interface and remains fully editable, if further adjustments are needed. Once final customizations are made, model training can be initiated with a single click. While the training proceeds, all relevant information, such as the training curves or the trained model, is periodically stored. Once trained, a model can be either fine-tuned on another dataset or used for prediction. The prediction process is straightforward for the end user (Supplementary Figure 2): the training configuration file is used to load the pretrained model, to normalize the new dataset, and to execute the prediction workflow. The local interface can also be employed in conjunction with a remote server, where computing resources are installed, and with OMERO^23^, a hosting platform for microscopy images (Supplementary Figure 3).

### A toolbox for bioimage analysts

Biom3d has been packaged as an easy-to-use Python library that can serve as an Application Programming Interface (API) and integrates a Command Line Interface (CLI). Exploiting the CLI enlarges Biom3d potentialities. For example, we have successfully run Biom3d on a High-Performance Computing (HPC) cluster managed by SLURM (Simple Linux Utility for Resource Management), leveraging job scheduling and resource allocation to efficiently process a wide variety of datasets. Both CLI and API can also help isolating individual processes such as data preprocessing, auto-configuration, or model training. Moreover, to fuse capabilities of multiple models trained on different datasets or with different configurations, Biom3d allows multi-model predictions. This last capability is particularly important to assemble the work of different teams across the world, working under different conditions. All these CLI options are directly accessible after installation of Biom3d Package, referenced in the Python Package Index (PyPI).

Python programmers may also be interested in exploiting the potential of the Configuration File and the eight types of modules. Configuration Files of Biom3d allows fine-tuning of more hyper-parameter settings more precisely than is feasible through the GUI alone, without the need to delve into Biom3d code. Biom3d modularity allows rapid experimentation with different configurations to identify the optimal setup for specific tasks or datasets. For example, the default deep learning model of Biom3d, which corresponds to the dynamic 3D U-Net available in nnU-Net framework, can easily be changed for another one. As a result, we demonstrated that a 3D EfficientU-Net model can outperform the 3D U-Net on the Pancreas dataset (Figure 3). Additionally, Biom3d is compatible with MONAI^10^, a public library integrating the latest biomedical models for multidimensional images, which enable quick use cutting-edge models.

### A framework for developers

Biom3d has been meticulously crafted with a focus on code clarity, enabling deep learning programmers familiar with libraries like PyTorch or TensorFlow to easily understand, modify, remove, or add components. It is also worth noting that Biom3d plays an important role in the process of comparing old and new code segments when creating new code based on existing works. This development strategy was, for instance, exploited to entirely redesign the original Dataset Module of nnU-Net. As Biom3d easily accepts alien code incorporation, the entire and original “Dataset Module” of nnU-Net was integrated as a Biom3d Module. Following this integration, a series of intermediate and hybrid Dataset Modules were developed and tested. Comparisons between old and new Dataset Modules could easily be made by editing a single line in the Configuration File. The weak points of the prototype Dataset Module were rapidly spotted and improved. The final Dataset Module, which relies on the TorchIO library^24^, is likely the reason why Biom3d outperforms nnU-Net on multiple tasks (Figure 3). This development strategy can be followed for all Biom3d Modules. Last, as each piece of Biom3d code can be isolated from the others, they can easily be extracted and reused for another independent project. This is particularly true not only for Biom3d Modules but even for individual functions such as auto-configuration or the image loading and saving.

## Discussion

Biom3d is a high-performing, and flexible set of tools applicable to various 3D segmentation problems, with numerous biological and medical applications. It surpasses in accuracy specialized tools, such as NucleusJ, while demonstrating state-of-the-art results on a wide variety of original biological questions, such as the quantification of the heterochromatin fraction in plants or the aorta lamella segmentation in rat. It responds to the needs of end users through easy to install and easy to use interfaces, enabling auto-configuration of the deep learning model training, and compatible with several image formats and with OMERO software^23^. To make Biom3d an even bigger part of the landscape of tools now available to biologist users, it will be made compatible with Napari^25^, ZeroCostDL4Mic^13,26^ and DeepImageJ^8^. In line with FAIR principles for AI^27^, future models will then be shared online in Open Neural Network Exchange (ONNX)^28^ format, on website such as HuggingFace.co or Bioimage.io, and future developments streamlined on MLflow (https://mlflow.org). To broaden Biom3d’s spectrum of applications, further developments should include instance segmentation, object tracking, image denoising as well as processing images in proprietary formats or with large N-dimension such as light sheet images and to offer the possibility of using images stored with the next generation file formats (NGFF) such as OME-Zarr^29^.

The implementation of all these developments will be facilitated by the modular code architecture of Biom3d. Biom3d has been designed as a sandbox, within which developers are strongly encouraged to integrate new code elements. It is coded in the Python programming language and based primarily on the Pytorch^30^ deep learning framework. This language and framework were chosen for their current popularity in the field of deep learning, so code implemented with them, such as MONAI, can be added to Biom3d with little to no effort. These constraints may yet pose some integration challenges for developers using other programming languages or deep learning libraries, such as TensorFlow^31^, with which StarDist was created, or JAX^32^, an emerging framework. While being highly modular, Biom3d code remains coupled to some extent. This occurs because it is needed in some instances to improve efficiency, by using shared internal functions across different components. It is minimized however to keep the advantages of modularity as interconnection can make the codebase more sensitive to changes as alterations to one section of the code might unintentionally impact other areas, complicating maintenance, and updates. The development of Biom3d represents a significant advance in the modularity of deep learning code, while maintaining performance and avoiding code redundancy. While further increases in modularity are possible, they may necessitate a significant additional investment in time and resources.

Finally, through the benchmarking performed on nucleus images in this article, we have pinpointed one of the main remaining limitations of deep learning methods: they depend on large, manually annotated datasets. While conventional image analysis tools can help provide partial annotations, manual interventions are inevitable, which also impedes the spread of deep learning methods. The modular code of Biom3d could be exploited to reduce this time-consuming process, by including methods such as active learning^33^, generative methods^34^, weakly supervised learning^35^, and self-supervised learning^36,37^. Biom3d has, for instance, successfully performed self-supervision for 3D images, which involved both image classification and image segmentation.

Most importantly, we hope that the intuitive design of Biom3d, from its graphical interfaces to its modular code architecture, will foster collaboration among diverse communities, including biologists, radiologists, microscopists, image analysts and developers, and will render it both reusable and sustainable.

## Online content

Biom3d is available at https://github.com/GuillaumeMougeot/Biom3d and as a notebook at https://colab.research.google.com/github/GuillaumeMougeot/Biom3d/blob/master/docs/Biom3d_colab.ipynb.

## Methods

### Requirements

Biom3d is a Python3 package essentially relying on numpy and Pytorch packages. Image reading and saving is ensured via scikit-image, SimpleITK and tifffile. Deep learning computation can be either executed on a CPU or a GPU. GPU execution requires the installation of CUDA and CuDNN libraries compatible with the Pytorch library as well as a GPU having at least 10 Gb of VRAM for training and 4 Gb for prediction.

### Graphical User Interfaces

The online interface is based on Google Colab and thus does not require accessing a local GPU. Its use requires the connection to a Google account. The training and prediction datasets must be uploaded to a Google Drive accessible via the interface. The code of the online interface is hidden for ergonomic reasons, but is accessible, and it has been made as simple as possible, exclusively using Biom3d framework. The online interface allows to preprocess a training dataset, to auto-configure a training process, to execute the training and to make prediction with a trained model. To take advantage of Biom3d modularity, the Configuration file is directly editable in Google Colab. While requiring access to a computing GPU, the local interface allows a higher degree of flexibility (Supplementary Figure 1 and Supplementary Figure 2). The user can start training or predicting on a local computer or on a private remote Linux server. In both cases, the interface is dynamic depending on user choices. If the user has a trained model and intends to fine-tune it, the interface displays a field for the path of the trained model and a field for the new dataset. If the user has a preprocessed dataset, only the configuration path will be asked. During prediction, if the user has access to an OMERO server (Supplementary Figure 3), the interface will ask for the user credentials and the OMERO dataset ID number. The OMERO dataset is then downloaded on the computing server, used for prediction, and the resulting images are eventually uploaded as a new OMERO dataset. For the remote interface, the user is additionally asked to provide a name to the training dataset which will automatically be sent to the computing server. After prediction, resulting images can either be downloaded locally or sent to an OMERO server.

### Preprocessing

Preprocessing consists in adapting a dataset to the training workflow and, inversely, adapting the training workflow to the dataset. The default preprocessing methodology has been designed and tested for 3D image segmentation. In the following explanation, masks are manually segmented images. The default preprocessing in Biom3d is a four-step process: (1) data reading and scanning to extract the data characteristics, (2) auto-configuration to create a Configuration file adapted to the data characteristics, (3) data normalization to uniformize image and mask dimensions and intensities, and (4) data splitting to separate training and validation sets. Preprocessing is illustrated in Supplementary Figure 4.

#### Data scanning

For any processing requiring image reading (data-scanning, data-loading, preprocessing, etc.), Biom3d adaptively reads 3D images stored in NumPy format (.npy), in TIFF format (.tif or .tiff) or in any other medical formats read by SimpleITK library (see list of available format on https://simpleitk.readthedocs.io/en/master/IO.html), such as Nifti format. After preprocessing or prediction, Biom3d saves 3D images or masks along with necessary meta-data. While compressed formats are appropriate for long term storage (such as Nifti or TIFF format), fast reading formats are preferred for deep learning applications (such as NumPy format). After preprocessing, training images and masks are thus stored in NumPy format. After prediction, Biom3d stores the resulting masks using the input image format and meta-data. As TIFF tagging system is unrestrictive regarding meta-data formatting, Biom3d image reader might not properly read meta-data originating from some proprietary formats, except if TIFF images have successfully been processed with Bio-format. The current image reader of Biom3d does not support proprietary formats, such as CZI. Once read, Biom3d extracts the dataset fingerprint from the images. This fingerprint includes the median image size, the mean and standard deviation of the image voxel intensities within the mask, the 5% and 95% percentiles of these intensities, and, if available, the median sampling, which represents the voxel dimensions in meters.

#### Auto-configuration

The auto-configuration process of Biom3d mainly consists in finding the best patch size and batch size to provide to the deep learning model, as well as the number and dimensions of the successive pooling layers occurring in the U-Net model. Biom3d heuristics produce similar results as nnU-Net ones, yet they have been simplified. Biom3d auto-configuration process only uses the median size of the input images. The ideal patch size is set to respect the median size proportions, while the product of its dimensions is smaller than a maximum patch size, by default defined as 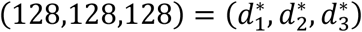, a value chosen for being adapted to GPUs with less than 10 Gb of VRAM. More formally, the goal is to find the exponents *α*_*i*_ in 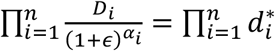,where *n* = 3 is the number of dimensions, 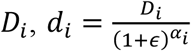 and 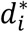 are the *i*-th dimension of the median size, the patch size and the maximum patch size respectively, and *ϵ* is a small number typically equal to 10^−3^. If all exponents *α*_*i*_ were assumed to be identical, their ideal value is 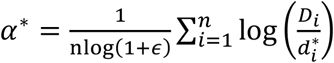. This value would ensure the preservation of the median size proportions. Yet, for large and strongly anisotropic images, this may cause some dimensions of the ideal patch size to be set smaller than 1. To prevent this, an upper bound *u*_*i*_ on *α*_*i*_’s values is set to be 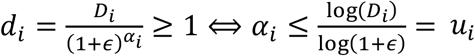. If *S* = {*i* ∈ ⟦1, *n*⟧, *α* > *u*_*i*_} ≠ ∅, then ∀*i* ∈ *S, α* = *u*_*i*_, and 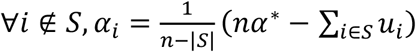, so to preserve ∑ *α*_*i*_ = *nα*\*. This process is reiterated until *S*= ∅. The final values of *α*_*i*_ can then be used to obtain 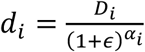, the patch size. For each dimension, the number of the 2-pooling in the U-Net model is then defined by 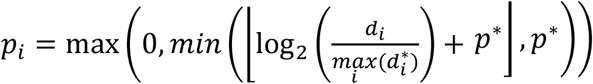, where *p*\* is the user-defined maximum number of 2-pooling, by default set to 5. To prevent features maps having non-integer dimensions after dimension reduction during 2-pooling, the patch size dimensions *d*_*i*_ are readjusted one last time to the closest multiple of 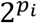. The batch is set to 2 and eventually increased if the GPU VRAM allows it. The auto-configuration results are stored in a new Configuration file in Python format, based on the default Configuration file included in Biom3d.

#### Data normalization

Data normalization encompasses image reshaping and resizing and voxel intensity normalizing (Supplementary Figure 5). Images are automatically reshaped to conform to the standard (channel, height, width, depth) dimensions. Biom3d accepts a wide range of dimension variants, even within the same dataset. If not specified by the user, Biom3d automatically detects the location of the channel dimension in 4-dimensional images. For each mask, a series of check-ups are performed to automatically spot and, eventually, correct annotation mistakes. For instance, within the same dataset, it can be found manual annotations with an inconsistent number or order of dimensions. If annotating with levels of grey, users could also mistakenly decide to use inconsistent levels of grey or more levels of grey than existing classes of objects. Depending on the mistake, Biom3d either corrects or warns the user. For instance, if two classes of object are expected and more than two were found in the mask, Biom3d automatically applies a threshold by considering as background the most recurrent class. In case where three or more classes are expected and even more classes are found in the mask, Biom3d displays an error. Biom3d is compatible with masks having 4 or 3 dimensions. Image intensities are then Z-normalized using either the mean and standard deviation of the image voxel intensities, or, if available, using the median mean and standard deviation of the dataset voxel intensities retrieved during data scanning. If the median sampling could be retrieved, images and masks are resized to all match the media sampling. Images are resized with trilinear interpolation while masks with nearest neighborhood. Finally, for each object class in the mask, a random sampling is performed to extract some foreground locations. These values will be used during data loading to rapidly locate foreground regions during image patch cropping. Once preprocessed, the images, masks and foreground (object of interest) locations are stored in automatically created output folders. The output folder paths are added to the Configuration file.

#### Data splitting

Following the cross-validation strategy, the dataset is by default split into 5 subsets called folds (Supplementary Figure 5). More formally, each image filename is associated with a random integer between 0 and 4. During training, if fold 0 is selected, images associated with 0 will be considered as validation images while remaining images will be considered as training images. If less than 10 images are present in the dataset, Biom3d reduces the number of folds to 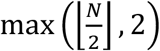, where N is the number of images in the dataset. If only one image is found in the dataset, Biom3d splits the image and mask in two along the largest image dimension – 80% of the image will be used for training and 20% for validation. Filenames with associated fold indices are stored in a CSV file that will be loaded during training. The CSV file path is added to the Configuration file.

### Data loading

Data loading consists in loading a batch of preprocessed data into computer memory and eventually performing data augmentation, for it to be prepared for the training workflow (Supplementary Figure 6). Several data loading modules for 3D image segmentation are available in Biom3d: one based on nnU-Net batchgenerators package, one based on TorchIO SubjectsDataset, and one based on Pytorch Dataset. The latter one is the default and will be detailed here (Supplementary Figure 6). The data loading module initialization first loads the image and mask filenames of training and validation sets using the CSV file created during data splitting. To fasten data loading, images and masks can be, on demand, loaded into computer memory. The initialization then creates the data augmentation transformations used during training with TorchIO package: random cropping, random affine transform, random anisotropy, random flipping, random intensity variation, random blurring, random noise, random patch swapping, and random contrast variation. The axes of the rotation in the random affine transform and of the random anisotropy depends on the patch size anisotropy. By default, anisotropy transforms, and rotation transforms are applied to every axis. If any patch size dimensions are bigger than three times the smallest patch size dimension, then only these dimensions are considered as valid axes to apply random anisotropy. In this scenario, only the smallest dimension is considered a valid axis for rotation transforms. Once this initialization is completed, the data loader is ready to use. During training data loading, a batch of foreground locations, images and masks are loaded into computer memory. The image and mask are then randomly cropped using the patch size. To do so, several constraints are considered. First, if rotation transforms are applied, the patch size is temporally enlarged beforehand to avoid empty corners appearing in images and masks once rotated. The enlarged patch size is set to be cubic and its dimensions equal to the largest diagonal of the patch. Second, foreground locations are used to crop one image and mask pair out of three in the batch. Among the list of possible foreground locations, one is selected to be the location of the center of the patch. Third, if the cropped image and mask are smaller than the patch size, then they are padded with zeros equally in all three dimensions. Once cropped, the images and masks are transformed with the rest of the data augmentations before being output. During validation data loading, only the random cropping is performed without any other form of data augmentation.

### Dynamic models

Biom3d includes several deep learning model definitions. The default deep learning model is a standard 3D U-Net model for semantic segmentation^1^. It has been implemented in a modular fashion, meaning that the encoder, the decoder and the convolutional blocks can work independently. More specifically, the default VGG encoder^2^ can straightforwardly serve as an independent classification model or can be replaced by any other encoder in the 3D U-Net model, such as the 3D EfficientNet^3^ included in Biom3d, the MONAI models, or any Pytorch model compatible with 3D images. The default 3D U-Net model architecture is dynamic and depends on the number *p*_*i*_ of successive pooling layers found for each dimension during the auto-configuration. The maximum number of 2-pooling along a given dimension *k* is determined by 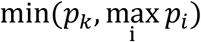.For each case where 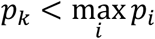,there will be 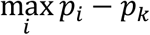 1-pooling layers, evenly distributed between the head and tail of the encoder. For instance, if the auto-configuration sets the ideal number of successive pooling layers to be (3,5,5), the pooling layers of the 3D U-Net will have the following list of kernel dimensions: ((1,2,2), (2,2,2), (2,2,2), (2,2,2), (1,2,2)). The definition of the rest of the model layers follows the original U-Net architecture.

### Training

Losses, metrics, callbacks as well as training and validation routines are all independent modules in Biom3d, the default ones being designed for 3D semantic segmentation. The default training loss is the sum of the class Dice score and the cross-entropy between the predicted and annotated masks. Default validation metrics include the Dice score and the intersection over union between the predicted and annotated masks. Even if the default behavior of these metrics requires the 3D annotated masks to be formatted such that each pixel value of the mask represents a single object class (0, 1, 2, etc.), Biom3d metrics are also compatible with 4D annotated masks formatted with an additional channel dimension representing each individual object class. In such setup, Biom3d metrics accept input pixel to be associated with multiple object classes. The default optimizer is the stochastic gradient descent with a Nesterov momentum of 0.99 and a weight decay of 3e-5. The default training routine uses the data loader to get a batch of data, passes it to the model then to the loss, computes the gradients and clips their norms, before updating the model parameters using the optimizer. The training routine periodically calls the callbacks (Supplementary Figure 7) to update the learning rate with cosine annealing, to print and store information about the training and validation, and to store the model parameters of the best performing model. Training and validation stored information includes the training and validation losses per epoch and prediction snapshot on the validation set. After loading the Configuration file, the Builder oversees the instantiation of the losses, metrics, callbacks and optimizer before executing the training and validation routines. This execution can be done using mixed precision and in parallel on multiple GPUs. For reproducibility reasons, once the training is finished, the output folder includes the Configuration file and the data splitting file, in addition to all the other information stored by the training callbacks. If interrupted, training can thus be restarted by solely using this output folder. Biom3d also allows to perform model finetuning or retraining by instantiating a new Builder using both an output folder and a new Configuration file.

### Predicting

Predicting with Biom3d encompasses three steps: (1) pre-processing, by default including image reading and normalization, (2) predicting, by default involving tiling the images and passing the tiles to the model, and (3) post-processing, by default implying discretizing model outputs, removing noise and saving the predicted masks. Each of these steps is an isolated Biom3d module. The modularity of Biom3d allows to reuse the exact same function for pre-processing as the one used to normalize training data. Before being passed to the deep learning model, input images are tiled with TorchIO grid sampler. For one input image, this function creates a series of patches equally distributed and overlapping by an overlap equal to half of the patch dimensions. To respect these two tiling constraints, the input image is eventually padded with zeros. Batches of patches are constituted and given to the model. To increase the prediction accuracy, each batch is augmented by flipping along the seven possible combinations of (x, y, z)-axis. Augmented predictions are then flipped back before being averaged. Predicted tiles are then aggregated using Hann filtering to reduce edge artefacts in patch overlapping regions. As data pre-processing involves data resampling, the aggregated prediction is finally resized back to the original image dimensions. As Biom3d framework is compatible with ensemble learning, it is also possible to aggregate predictions coming from different models. If such case, model outputs are simply averaged before being post-processed. Post-processing then starts by discretizing the prediction. For 4D masks, discretizing means to apply a threshold of 0.5 to the output of the sigmoid function applied to the model output. For 3D masks, discretizing means to retrieve the argmax of the output of the SoftMax function applied to the model output. On user request, two distinct noise removal strategies can then be applied to remove too small, segmented regions. Connected components can thus be computed to retrieve either the biggest segmented object or objects which volumes are higher than an Otsu threshold determined using the volume distribution of all connected components. Finally, the post-processed prediction is automatically saved along with input image metadata.

### Evaluation

Controlling the quality of a trained model can be done with Biom3d by either using the local GUI or the API. For 3D segmentation, one folder containing 3D ground truth masks of images different from the training set and another folder with the corresponding predictions can be passed to Biom3d to retrieve the average Dice score on this set. To speed up experiments and benchmarking, Biom3d also includes Python scripts that allow the preprocessing, training, prediction, and evaluation of a new deep learning model on a new dataset to be executed with a single command.

## Data availability

All datasets and trained models are described in the Supplementary Table 1, including links to see and download them. Custom datasets are hosted on the OMERO repository of the Mésocentre Clermont Auvergne (see Supplementary Table 1).

## Code availability

Biom3d is a public Python package referenced in the Python Package Index (https://pypi.org/project/biom3d/). The latest version of the code and documentation of Biom3d can be found on GitHub (https://github.com/GuillaumeMougeot/biom3d).

## Acknowledgements

We would like to thank Adama Nana for his support with benchmarking nucleus segmentation methods, the Auvergne Bioinformatic (AuBi) platform and the Mésocentre Clermont-Auvergne of the Université Clermont Auvergne for providing help, computing and storage resources facilities and the iGReD CLIC microscopy facilities for image acquisition. This work was partially achieved using HPC resources from GENCI–IDRIS (Grant 2022-AD011013709) on the supercomputer Jean Zay’s A100 partition. We thank Dr. Fredy Barneche (CNRS, IPBS Sorbonne) for providing us with seeds; Xiaowen Liang (Université de Reims Champagne Ardenne) for providing us with annotated images of rat aorta; Cynthia Dennis (iGReD) for providing us with an annotated dataset of epithelial cells of drosophila ovaries and Nicolas Allègre (iGReD) for stained mouse embryos. Students in the Master 1 Bioinformatics program (UCA) have contributed to the annotation of nuclei images and the drafting of documentation for biologists (graduating classes 2023-2024 and 2025).

This work was supported by Agence Nationale de la Recherche of the French government through the programme ‘Investissements d’Avenir’ (16-IDEX-0001 CAP 20-25), ‘Seed Chrom’ ANR-22-CE20-0028; Oxford Brookes University, Université Clermont Auvergne, Centre National de la Recherche Scientifique, Institut National de la Recherche et de la Santé, the Université Reims Champagne Ardenne, the European Regional Development Fund (FEDER), EMERGENCE (16-IDEX-0001 CAP 20-25) and the trainee grants funding GDR IMABIO (France). A.P., C.T., D.E.E., K.G. and S.D. are part of the International Plant Nucleus Consortium (IPNC; https://radar.brookes.ac.uk). The authors acknowledge Synchrotron SOLEIL to have provided beamtime for Figure 2 under project no. 20211303. ANATOMIX is an Equipment of Excellence (EQUIPEX) funded by the Investments for the Future program of the French National Research Agency (ANR), project NanoimagesX, grant no. ANR-11-EQPX-0031.

## Author contributions

G.M. conceptualized the coding philosophy and undertook most of the programming work and experiments. D.E.E., F.C., C.T., E.P. and S.D. provided ideas for the project and obtained funding. S.S. helped debugging and improving the user interface. G.M. and S.D. wrote the manuscript with input from C.T., F.C., D.E.E. and E.P. G.M. and S.D. performed data analysis and prepared the figures. A.P., H.A., P.P., S.D., S.A. and N.F. provided annotated images and were beta testers of Biom3d.

## Competing interests

The authors declare no competing or financial interests.

**Supplementary Figure 1.**
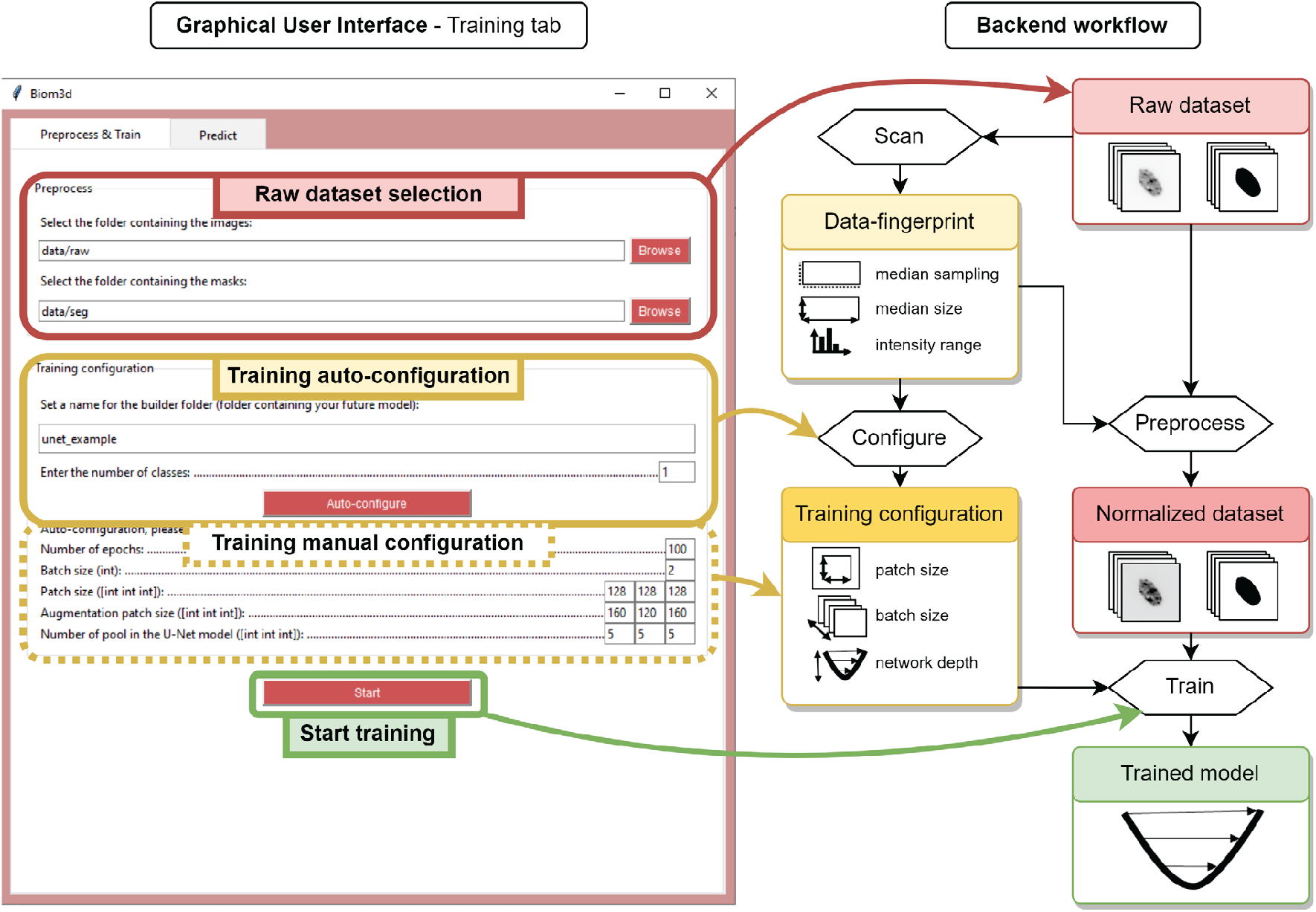
Training workflow of Biom3d in the local graphical interface. (*Left*) Training tab of the local graphical interface. The user specifies the path to the folders containing training images and annotations, then defines a name for the Configuration file and the future trained model, and the auto-configuration can start. Automatically defined parameters can be adjusted manually, if needed, before starting the training. **(*Right*) Backend workflow**. Once the “Auto-configuration” button is pressed, the data-preprocessing starts. The dataset key elements (median shape etc.) are extracted and used to normalize all the images and to define training configuration (patch size etc.). If the “Start” button is pressed, a deep learning model will be trained and saved along with the pre-processing methodology (data-fingerprint).

**Supplementary Figure 2.**
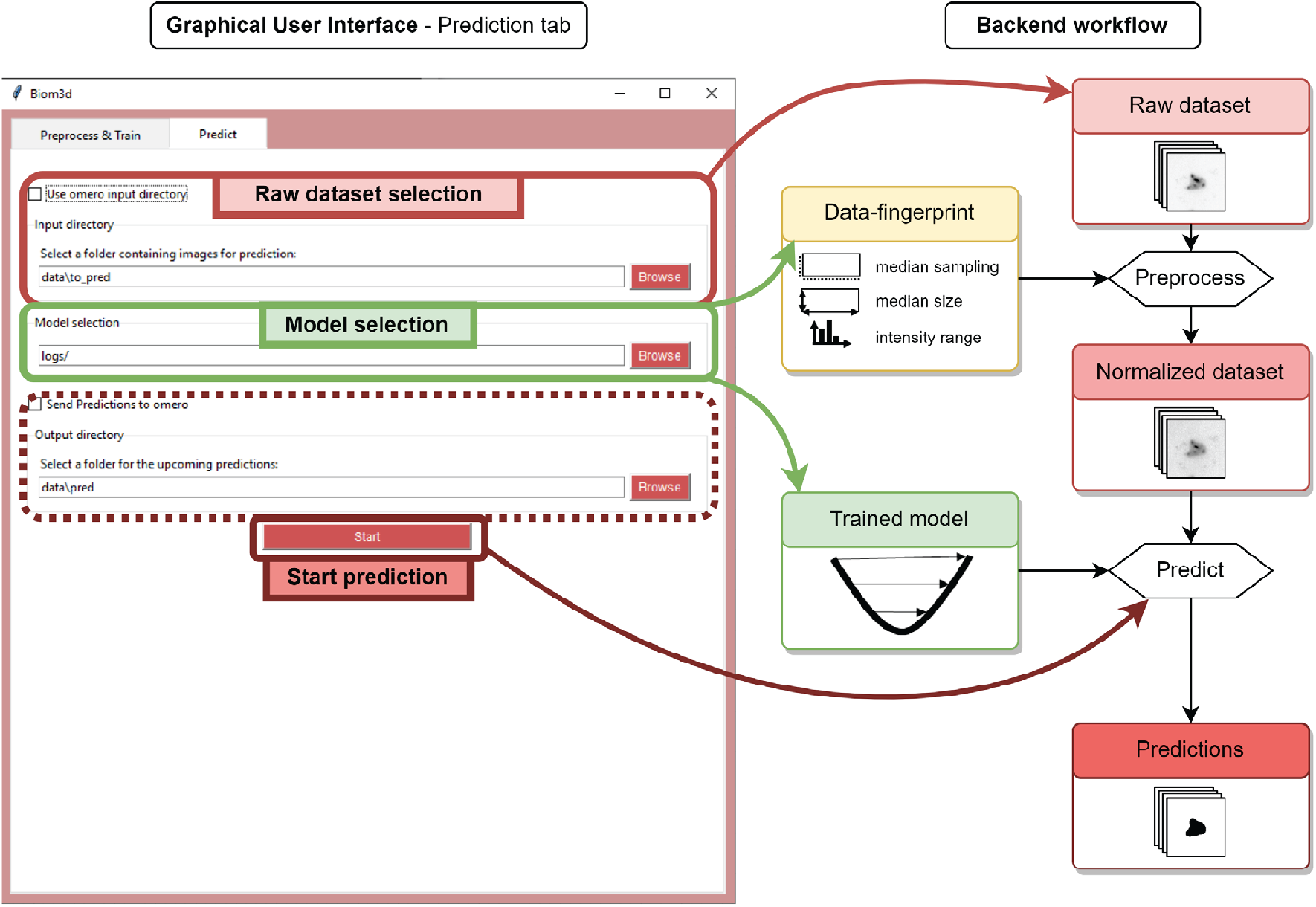
Prediction workflow. (*Left*) Prediction tab of the graphical interface. The user chooses a folder containing raw images, the path to a trained model and a folder for the future predictions. The predictions start when the “Start” button is pressed. **(*Right*) Backend workflow**. The raw images are normalized using the data-fingerprint of the training dataset. The trained model is then loaded and used to compute predictions.

**Supplementary Figure 3.**
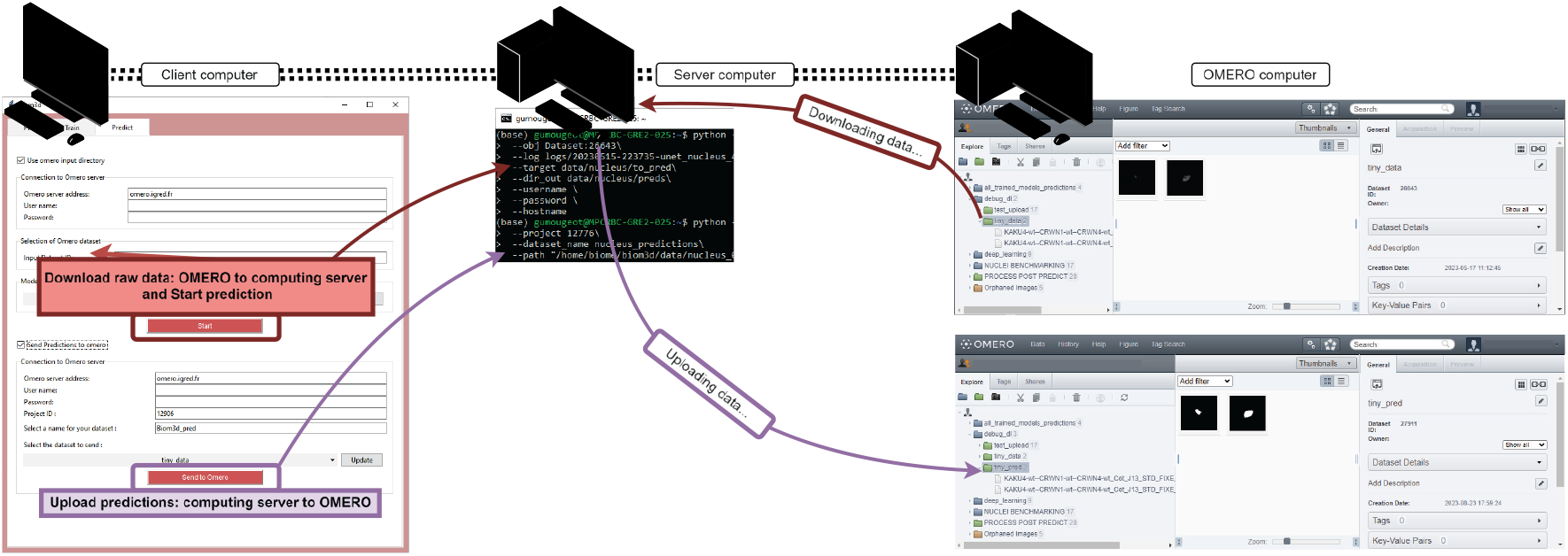
Remote access and OMERO access to Biom3d. Remote access and OMERO access can work together or independently. A combined use case is represented. The local interface (*left*) allows the access to a remote server remote (*middle*) running Biom3d training or prediction workflows. If used in combination with an OMERO server (right), the remote server will download raw images from an OMERO dataset and upload them back in a new OMERO dataset.

**Supplementary Figure 4.**
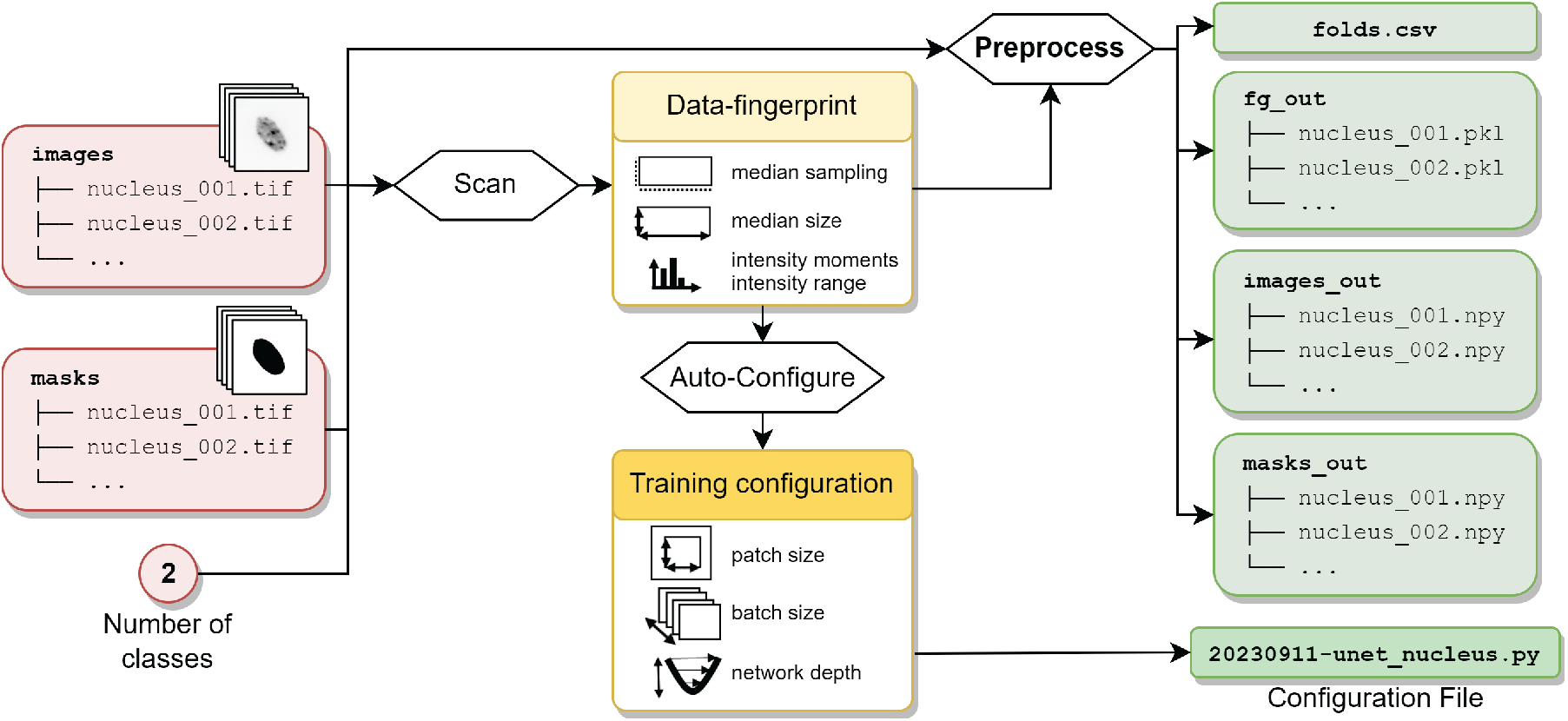
Auto-configuration and Preprocessing of training dataset for 3D segmentation. Image and mask folders (*red, left*) are scanned to extract their data-fingerprint. The data-fingerprint is used to preprocess the images and masks and to automatically configure the future training (*yellow, centre*). The outputs of these two steps (*green, right*) are: a CSV file (*folds*.*csv*) describing which files will be used for training or validation, a Configuration file, the pre-processed images (*images_out*) and masks (*masks_out*), and the location of the foreground voxels (*fg_out*).

**Supplementary Figure 5.**
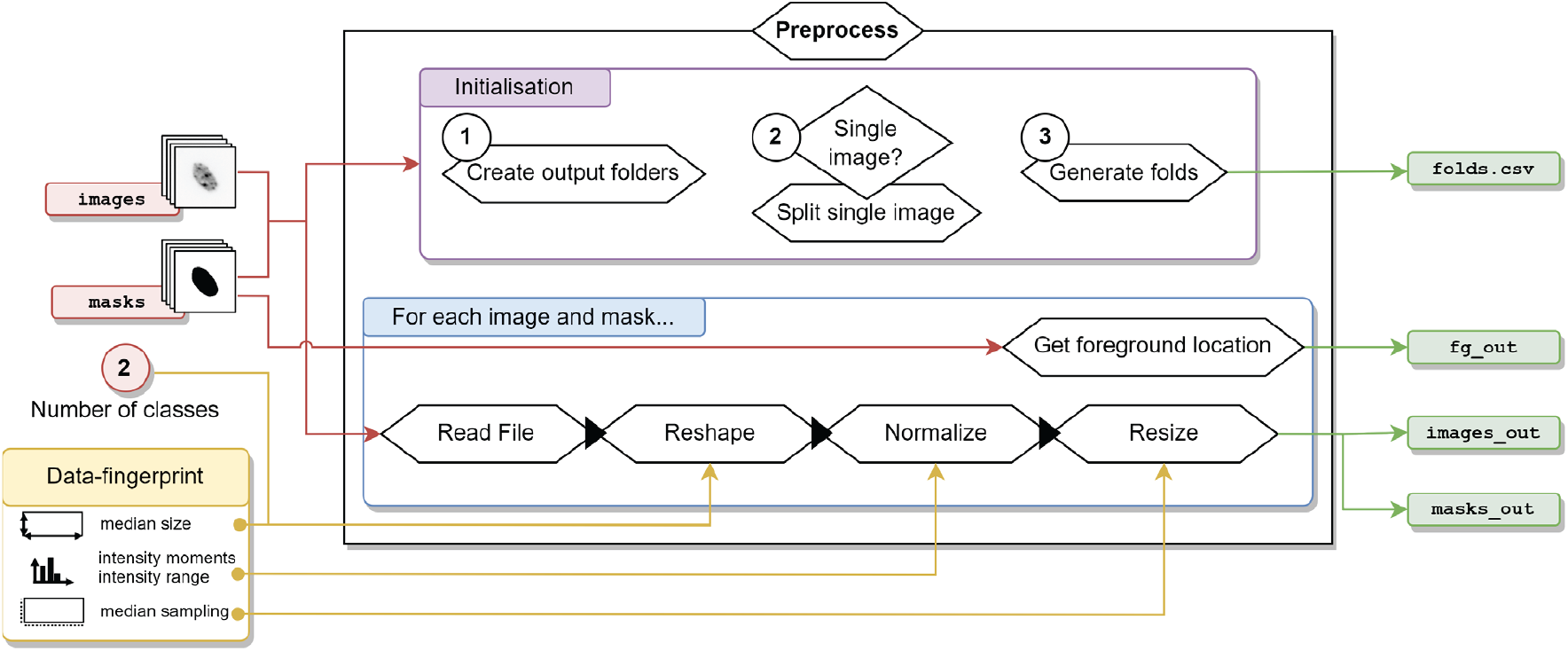
Training data normalization and splitting for 3D segmentation. The preprocessing starts (*purple box, initialization*) by (1) creating the three output folders (*fg_out, images_out, masks_out*), (2) optionally splitting single image/mask and (3) splitting the dataset into training and validation folds (*folds*.*csv*). Afterwards, each image and mask (*blue box*) are read (*Read File*), independently of their format, reshaped (*Reshape*) so to have exactly 4 dimensions in (*channel, depth, heigh, width*) format, z-normalized for images and uniformized for masks (*Normalize*), and resized (*Resize*) so all images and masks have the same sampling. Finally, the locations of foreground voxels are extracted and stored in the appropriate output folder (*fg_out*).

**Supplementary Figure 6.**
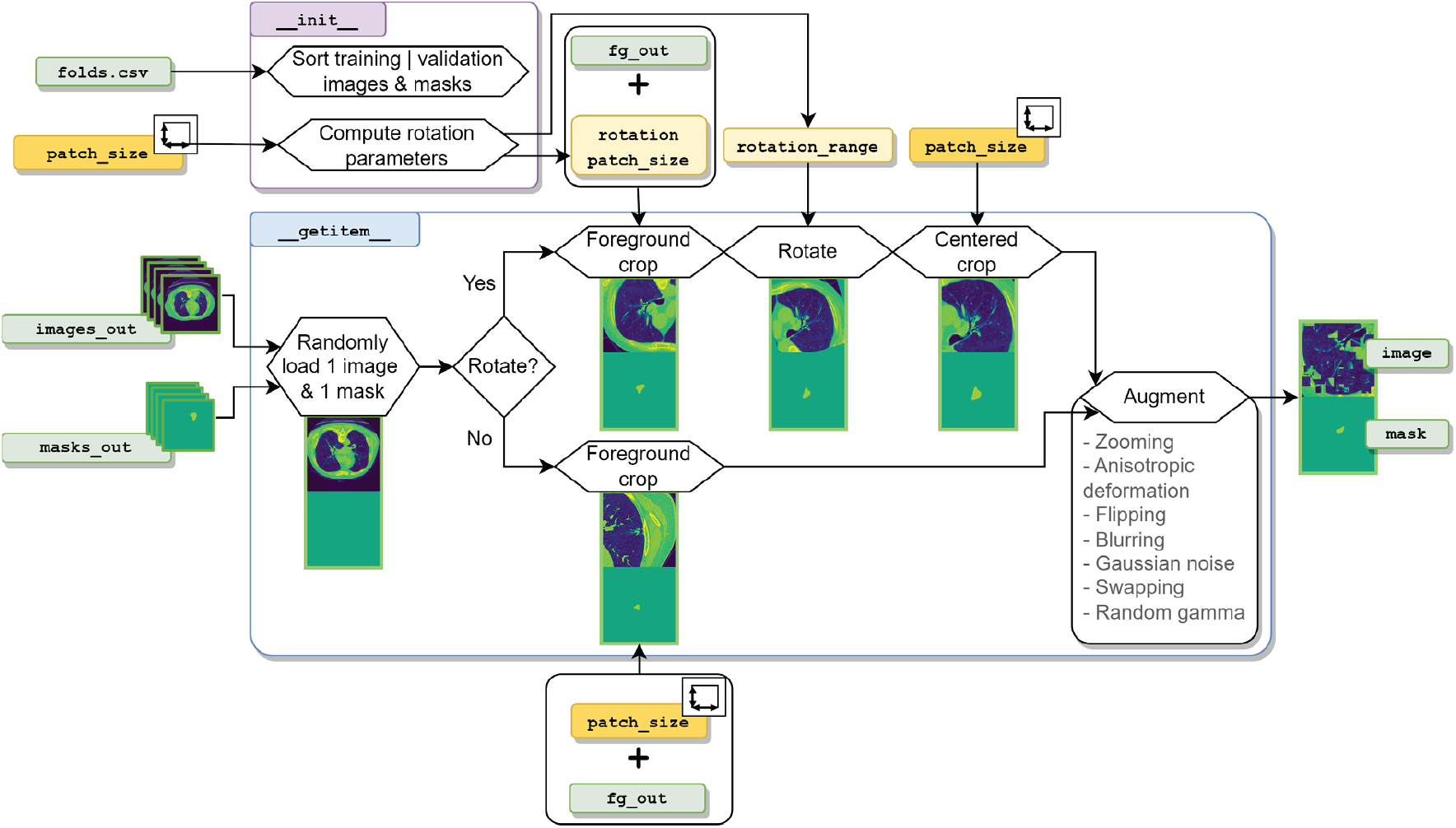
Biom3d default Dataset Module for 3D segmentation. In the `_init_` class function (*purple*), the CSV file is used to sort training images from validation images and the patch size is used to determine the parameters of the rotation transformation (rotation angle and rotation patch size, *yellow*). In the `_getitem_` class function (*blue*), one image and one mask are loaded into computer memory from their local folder. Those are then cropped in regions where foreground objects are located. If rotation augmentation is active, then the foreground crop is performed with a larger patch size before being cropped a second time to discard unwanted empty regions in the image corners. Another series of augmentations is finally applied to obtain a ready to use pair of image and mask patches.

**Supplementary Figure 7.**
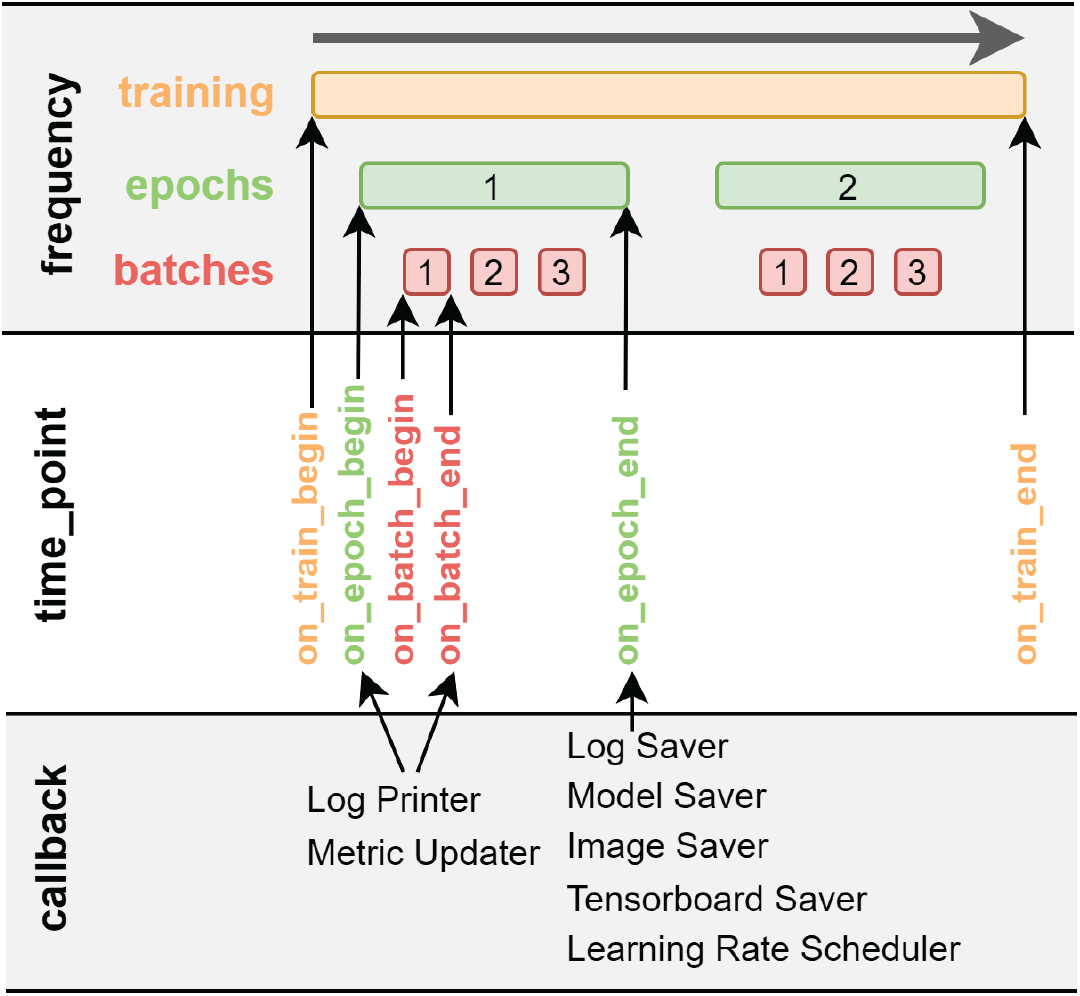
Callback Module principle. The whole training (top row, orange) is divided into epochs (green) themselves divided into batches (red). Callbacks are Python classes that can have one or more class-functions, each representing one of 6 different time points (middle row). Biom3d currently has 7 types of Callback Modules (bottom row).

